# Visual deprivation in adulthood engages presynaptic plasticity of thalamocortical synapses

**DOI:** 10.64898/2026.08.17.745313

**Authors:** Sachiko Murase, Daniel Severin, Andrew Pranger, Louis Dye, Cristian Moreno, Lukas Mesik, Ji Liu, Alfredo Kirkwood, Elizabeth M. Quinlan

**Affiliations:** Department of Biology, University of Maryland, College Park, MD 20742; Mind/Brain Institute, Johns Hopkins University, Baltimore MD 21218; Department of Neuroscience, John Hopkins University, Baltimore, MD 21205; Department of Neuroscience, School of Medicine and Public Health University of Wisconsin, Madison WI 53507; NICHD Eunice Kennedy Shriver, Microscopy & Imaging Core, NIH, Bethesda, MD 20892

## Abstract

Synaptic plasticity between neurons in first order thalamus and layer 4 cortex is greatest during an early postnatal critical period and was thought to decrease irreversibly with age. However, here we show that robust and reversible plasticity can be induced in adult mice at synapses between dLGN axons and visual cortex layer 4 neurons by prolonged dark exposure (DE) and light reintroduction (LRx). Unexpectedly, the experience-dependent change in synaptic strength was mediated by a change in presynaptic structure, organization and function. DE/LRx declusters/clusters synaptic vesicles and reorganizes presynaptic molecular geometry. Furthermore, DE/LRx decreases/increases visually-evoked and spontaneous calcium signaling and neurotransmitter release probability in dLGN axonal boutons. The presynaptic plasticity mechanism described here has a high threshold for engagement which would allow maintenance of synapse stability across a wide activity range, and reactivation of thalamocortical plasticity in extraordinary conditions.

**Significance:** The age-dependent reduction in plasticity at thalamocortical synapses contributes to the developmental constraints on the reversal of amblyopia beyond a postnatal critical period. Here we describe that robust plasticity is engaged at thalamocortical synapses in adults by visual deprivation, a non-invasive manipulation with therapeutic potential.

## Introduction

Synaptic plasticity is constrained by age, endowing mature neuronal circuits with enhanced stability. However, we have previously shown robust receptive field plasticity can be induced in the adult rodent primary visual cortex (V1) by visual deprivation through prolonged dark exposure and subsequent light reintroduction (He et al., 2006, 2007). The ability to enhance plasticity in adults through manipulations of visual experience holds therapeutic value. Accordingly, in a rat model of severe amblyopia, visual stimulation following DE drives the recovery of vision in the amblyopic visual pathway (He et al., 2007; Montey and Quinlan, 2011; Eaton et al., 2016). The rejuvenation of plasticity by DE has been also demonstrated in felines and mice (Duffy et al., 2016; Stodieck et al., 2014; Murase et al., 2017; Jeon et al., 2022). However, little is known of the mechanisms that couple prolonged sensory deprivation with an enhancement of synaptic plasticity in sensory cortex.

Activity-dependent synaptic plasticity can either amplify or compensate for changes in neuronal activity. Visual deprivation induces a well-documented homeostatic increase in the strength of intracortical (IC) synapses in V1 (Wen and Turrigiano, 2024). DE increases the expression of the GluN2b NMDAR subunit in adult rodent V1 (He et al., 2006), which lowers the threshold for correlation-based synaptic plasticity (Cooper and Bear, 2012; Lee and Kirkwood, 2019) and increases spontaneous neuronal spiking. The homeostatic increase in mEPSC amplitudes in superficial lamina of V1 induced by visual deprivation is not restricted to a juvenile critical period (Goel et al., 2006; Goel and Lee, 2007). Brief dark exposure induces rapid homeostatic changes in the concentration of glutamate/glutamine in human V1 and amplifies the shift in ocular dominance in response to monocular deprivation (Min et al., 2023). In contrast, the expression of plasticity at synapses between first order sensory thalamus and cortex is thought to be constrained in early postnatal development. Classic experiments in slices of post-critical period rat barrel cortex demonstrated that EPSP slope/amplitude at thalamocortical synapses are unchanged following stimulation protocols for pairing-induced long-term potentiation (LTP), NMDAR-dependent long-term depression (LTD) and silent synapse conversion (Crair and Malenka, 1995; Isaac et al., 1997; Feldman et al., 1998). Similarly, in slices of primary visual cortex, paired stimulation of white matter and layer 4 is ineffective in inducing LTP after an early critical period (Jiang et al., 2007).

Nonetheless, our previous work suggested that DE and LRx could enhance plasticity at thalamocortical synapses (Montey and Quinlan, 2011; Murase et al., 2017). To ascertain if DE/LRx can modify the efficacy of feed-forward synapses from the dorsal lateral geniculate nucleus (dLGN) to the visual cortex in adult mice, we employed a series of orthogonal experimental approaches to examine the structure and function of these synapses. 1) Transmission electron micrographs revealed reversible declustering/clustering of synaptic vesicles in thalamic boutons following DE and LRx. 2) FRET-based intermolecular proximity analysis demonstrated declustering/clustering of a presynaptic cytomatrix protein in dLGN boutons. 3) Intravital calcium imaging revealed that DE decreased and LRx increased the magnitude of spontaneous and visually-evoked thalamic bouton calcium signals. 4) Optogenetic stimulation of dLGN boutons suggest that presynaptic release probability was decreased by DE and increased by LRx. Thus, latent plasticity with a presynaptic locus, can be unmasked by manipulation of visual experience.

## Materials and methods

### Subjects

C57BL/6J mice were purchased from the Jackson Laboratory (Bar Harbor, ME, USA). Equal numbers of adult (>P90) males and females were used. Mice were raised in a 12:12 hour (h) light:dark cycle. All procedures conformed to the guidelines of the University of Maryland, University of Wisconsin-Madison and Johns Hopkins University Institutional Animal Care and Use Committees. Control experiments were performed (or subjects were sacrificed) 6 h into the light phase.

### Electron Microscopy

Mice were transcardially perfused with 2.5% glutaraldehyde and 4% paraformaldehyde (PFA) in phosphate buffered saline (PBS, pH 7.4). Brains were post-fixed overnight and stored in 0.01% NaN_3_ in PBS at 4°C. Coronal sections (100 microns) were made using a Leica vibratome (VT1000) and V1b and the hippocampal commissure were isolated. Sections were rinsed in 0.1M sodium cacodylate buffer. The following processing steps were then carried out using a variable wattage Pelco BioWave Pro microwave oven (Ted Pella, Inc., Redding, CA): post-fixation in 1% osmium tetroxide made in 0.1 M sodium cacodylate buffer, rinse in double distilled water (DDW), 2% (aq.) uranyl acetate enhancement, DDW rinse, ethanol dehydration series up to 100% ethanol and propylene oxide, followed by an Embed-812 resin (Electron Microscopy Sciences, Hatfield, PA) infiltration series up to 100% resin. The epoxy resin was polymerized for 20 h in an oven at 60°C. Ultra-thin sections were cut on a Leica EM-UC7 Ultramicrotome (90 nm). Thin sections were placed on 200 mesh copper grids and post-stained with uranyl acetate and lead citrate. Images were collected 360±40 µm from dura. Images were acquired with a JEOL-1400 Transmission Electron Microscope operating at 80kV and an AMT BioSprint-29 camera at 25K times magnification at 0.37 nm/pixel resolution. All synaptic profiles with high contrast boundaries delineating the presynaptic membrane, active zone, postsynaptic membrane and postsynaptic density were quantified. Image acquisition and analysis were performed blind. For quantification of ultrastructural variables, we used ultrathin sections chosen randomly to avoid sampling bias and adopted the area of the presynaptic terminal as a proxy for presynaptic origin (Schoonover et al., 2014; Bopp et al., 2017).

### FRET-based molecular proximity assay

Subjects were anesthetized with 4% isoflurane in O_2_ and perfused with PBS followed by 4% PFA in PBS. The brains were post-fixed in 4% PFA for 24 h, followed by 30% sucrose for 24 h, and cryo-protectant solution for 24 h (0.58 M sucrose, 30% (v/v) ethylene glycol, 3 mM sodium azide, 0.64 M sodium phosphate, pH 7.4). Coronal sections (40 μm) were cut on a Leica vibratome (VT1000). Sections were blocked with 4% normal goat serum (NGS) containing 0.4% TritonX-100 and 0.1% Tween-20 in PBS for 30 min. The primary antibody was presented in blocking solution overnight at 4°C, followed by secondary antibodies for 6 hours.

The following antibodies/dilutions were used: Monoclonal mouse anti-Bassoon (Bsn, NeuroMab) RRID: AB_2895685, 1:1000; goat anti-mouse IgG Alexa-568 conjugated (Thermo Fisher Scientific) RRDI: AB_2534072, 1:1000; goat anti-mouse IgG Alexa-647 conjugated (Thermo Fisher Scientific) RRID: AB_2535804, 1:1000; unless the dilution is specified. FRET experiments were performed with the anti-Bsn monoclonal mouse antibody and a 50:50 mixture of the two secondary antibodies. Thalamic boutons were labeled with polyclonal guinea pig anti-vesicle glutamate transporter 2 (VGluT2, SynapticSystems) RRID: AB_887884, 1:1000, and visualized with goat anti-guinea pig IgG Alexa-488 conjugated (Thermo Fisher Scientific) RRID: AB_2534117, 1: 1000.

Images were acquired on a Leica SP5x confocal microscope with a 60X Oil lens (HCX PL APO CS, NA = 1.4) at Ex = 488 nm with detection range (500 – 530 nm) for VGluT2, and Ex = 543 nm, with detection range for Alexa-568 (560 – 615 nm) and Alexa-647 (650 – 720 nm) for Bsn FRET. VGluT2 and Bsn puncta were analyzed in single Z-section images in Fiji (NIH) following thresholding (auto threshold + 20) and size exclusion (0.2–2.0 μm^2^) using the *analyze particles* function. Image acquisition and analysis were performed blind.

### Intravital imaging of axonal GCaMP

Axonal GCaMP6s was delivered to dLGN (from Bregma AP: 2.1 mm, ML: 2.2 mm, DV: 2.5 mm) via AAV5-Ef1a-DIO-Synaptophysin-GCaMP6s (addgene, Cat# 105715-AAV5, 2.0 x 10^13^ GS/ml) in a Hamilton syringe attached to a Microsyringe Pump Controller (World Precision Instruments) at a rate of 100 nl/min (total volume of 750 nl) to adult (P>90) VGluT2-Cre mice (Jax strain 028863).

A cranial window consisting of two 3 mm diameter coverslips glued with optical adhesive (Norland 71, Edmund Optics) to a 5 mm diameter coverslip was implanted. The gap between the skull and glass was sealed with silicone elastomer (Kwik-Sil). Instant adhesive Loctite 454 (Henkel) was used to adhere an aluminum head post to the skull and to cover the exposed skull. Black dental cement (iron oxide powder, AlphaChemical mixed with white powder, Dentsply) was used to coat the surface to minimize light reflections. Subjects were imaged ≥ 3 weeks following surgery.

Awake subjects were placed in a holding tube and immobilized by a head post clamp. Prior to the first imaging session, the subjects were habituated to the holding tube at least twice for >30 min. A shield was placed around the gap between the cranial window and the objective lens to block light contamination from the visual stimulus during image acquisition. An Olympus FVMPE-RS Multiphoton Laser Scanning Microscope controlled by Fluoview software with a 25x NA 1.05 water immersion objective lens was used to acquire time lapse fluorescence images. An Insight X3-OL laser was tuned to 940 nm for imaging GCaMP (Ex max 490 nm). Fluorescence emission was detected through a dichroic mirror (495-540 nm). The field of view was 169.71 µm x 169.71 µm (512x512 pixels), 360±40 µm from the brain surface. Minimum laser power (59 mW) and PMT gain (400) necessary to image at this depth was used. We observed no photobleaching with these imaging parameters. Images of visually evoked GCaMP ΔF/F were acquired in resonant scanning mode at 28 Hz. Using the Suite2P package (Pachitariu et al., 2017), movement artifacts were corrected using the average intensity of the full image stack as a template. Region of interests (ROIs) were detected in axon/bouton mode (correction factor, 0.7). Calcium signals were deconvoluted using OASIS (Friedrich et al., 2017).

To evoke visual responses in boutons, subjects received monocular visual stimuli controlled by PsychToolBox plugin in MATLAB (random order of 5 repeats of 12 directions of square wave gratings drifting at 1 Hz at 0.05 cycle/degree (cpd), 100% contrast for 2.5 s interleaved with 2.5 s intensity-matched (28 cd/m^2^) grey scale, delivered by a 23" display (Acer LCD Monitor) 28 cm in front of the eyes). Calculation of ΔF/F = (F-F0)/F0, where F corresponds to the fluorescence intensity at a given time point and F0 corresponds to mean fluorescent intensity for 1 s before the visual stimulus onset. The inclusion criterion for visually responsive boutons was: 1) an evoked peak ΔF/F value (0.25 - 2.0 sec after onset of a visual stimulus in any direction) greater than 4X the standard deviation (STD) of pre-stimulus baseline (F_0_, −1.0 to 0 sec) for ≥ 40% trials and 2) a STD of ΔF/F during the rising phase of response (0.25 to 1.0 sec after visual stimulus onset) greater than 2 X STD of baseline.

Orientation selectivity and direction selectivity were calculated using the following equations: the direction vector, dirR, was defined as dirR = (ΣR(θ) x cosθ/ΣR(θ), ΣR(θ) x sinθ/ΣR(θ)), where R(θ) represents the response (ΔF/F) at the stimulation direction, θ. Direction selectivity was calculated using the direction circular variance, dirCV = 1 − |dirR| (Mazurek et al., 2014). The orientation vector, R was defined as R = (ΣR(θ) x cos2θ/ΣR(θ), ΣR(θ) x sin2θ/ΣR(θ)). The orientation selectivity was calculated using the orientation circular variance, CV = 1 − |R|, which is more robust for neurons with weak tuning (Mazurek et al., 2014). The area under the curve (AUC) of deconvolved spontaneous calcium signals was calculated in MATLAB for 1 s in the absence of visual stimulation.

### V1 slice preparation and optogenetic thalamocortical synapse isolation

AAV5-CaMKIIa-hChR_2_(H134R)-mCherry (addgene, Cat# 26975-AAV5, 1.4 x 10^13^ GS/mL, 300 nl) was delivered to adult C57BL/6J mouse dLGN (from Bregma AP: 2.3 mm, ML: 2.0 mm, DV: 2.4 mm) 3 weeks prior to recording at >postnatal day 90 (>P90). Mice were anesthetized using isoflurane vapors; after disappearance of the corneal reflex mice were transcardially perfused with ice-cold dissection buffer containing (in mM): 212.7 sucrose, 5 KCl, 1.25 NaH_2_PO_4_, 10 MgCl_2_, 0.5 CaCl_2_, 26 NaHCO_3_, and 10 dextrose, saturated with 95% O_2_/5% CO_2_ (pH 7.4). The brain was rapidly removed, immersed in ice-cold dissection buffer and sectioned (300 µm) using a vibratome (Leica VT1200S). Coronal slices containing primary visual cortex were transferred to artificial cerebrospinal fluid (ACSF), incubated at 30°C for 30 min, room temperature for 30 min, and then transferred to the recording chamber. ACSF was identical to dissection buffer except that sucrose was replaced by124 mM NaCl, MgCl_2_ was lowered to 1 mM, and CaCl_2_ was raised to 2 mM. Visualized whole-cell recordings were made from pyramidal neurons in layer 4 of V1 with glass pipettes (3-5 MΩ) filled with (in mM) 8 KCl, 125 cesium gluconate, 10 HEPES, 1 EGTA, 4 MgATP, 0.5NaGTP, and 5 QX-314 (pH 7.2 to 7.3 and 280 to 295 mOsm). Biocytin (1 mg/ml) was added to the internal solution for *post hoc* cell identification. ChR_2_-containing axon terminals were activated using a 470 nm wavelength LED (3 ms duration; Thorlabs) through a 40x objective lens. EPSCs were recorded in voltage–clamp at −70 mV.

Sr^2+^-desynchronized miniature EPSCs (Sr^2+^ mEPSCs) were evoked by the light intensity triggering a response in 50% of trials (half-maximal effect light intensity, 470 nm, 3 msec) following replacement of Ca^2+^ with 4 mM Sr^2+^ and raising Mg^2+^ concentration to 4 mM (Dodge Jr. et al., 1969). To ensure that light-evoked responses are monosynaptic, recordings were performed in the presence of 1 μM TTX and 100 μM 4-AP (Petreanu et al., 2009; Whitt et al., 2022). Only cells with series resistance <25 MΩ and <25% variation over the experiment were included. Data was filtered at 4 kHz and digitized at 10 kHz using Igor Pro (Wave Metrics).

To quantify the response to repetitive optogenetic stimulation, 15 light pulses were delivered at 25 Hz at the minimal light intensity to evoke a response in 100% of trials. EPSCs were recorded in the presence of 100 μM APV and 20 μM bicuculline. The shape of the EPSC was examined, and cells were excluded from analysis if the rising phase exhibited any inflection points, which could indicate a polysynaptic response. The paired pulse ratio was calculated as 2^nd^ EPSC/1^st^ EPSC. The normalized EPSC at steady state was calculated as the mean of 11^th^ to 15^th^ EPSC. The release probability and the recovery rate were calculated as described in Bridi et al., 2020 based on Wesseling and Lo, 2002. Briefly, equations (2) and (3) were derived from equation (1) as shown below:

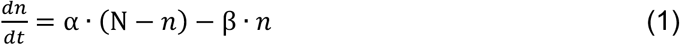

Where α denotes the recovery rate, β denotes the release probability, *n* represents the vesicles available for release, and *N* represents the total number of vesicles in the readily releasable pool.

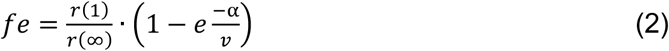

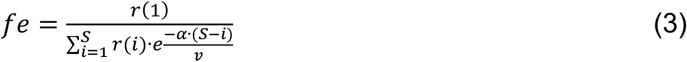

Where *fe* is the initial release probability, *r*(1) is the amount of neurotransmitter released by the first stimulus in the train, *r*(∞) represents the release at steady-state during the train, *v* is the stimulation frequency, *S* is the total number of stimuli in the train, *i* denotes the *i*-th stimulus, *r*(*i*) is the release corresponding to the *i*-th stimulus, and *r*(*i*)/*r*(1) represents the normalized amplitude of the *i*-th EPSC in the train. The values for *fe* and *α* were determined by solving equations (2) and (3).

### Biocytin processing

After recordings, slices were fixed in 4% paraformaldehyde overnight at 4°C. Slices were rinsed 2 × 10 min in 0.1 M phosphate buffer (PB) containing 1.9 mM NaH_2_PO_4_ · H_2_O and 8.1 mM Na_2_HPO_4_ at room temperature and permeabilized in 2% Triton X-100 in 0.01 mM PB for 1 h. Slices were then incubated in avidin-Alexa Fluor 633 conjugate diluted 1:2000 in 1% Triton X-100/0.01 M PB overnight at 4°C in the dark. After the incubation, slices were washed 2 × 10 min in 0.01 M PB, mounted on glass slides, and allowed to dry. Slides were cover-slipped with Prolong Gold Anti-fade (Invitrogen) mounting medium and sealed with nail polish. Images were taken using a Zeiss LSM 700 confocal microscope.

### Statistics

An unpaired two-tailed Student’s t-test was used to determine the significance between two independent experimental groups. A Mann-Whitney test was applied to EM analyses due to the bimodal distribution of presynaptic area. A one-way ANOVA was performed to determine the significance between three or more independent experimental groups. For spike train experiments, a two-way ANOVA was performed. Kolmogorov-Smirnov tests were used to compare the distributions of AUC.

## Results

The distinct molecular geometry and morphology of thalamocortical and intracortical synapses was used to determine synapse subtype-specific changes in transmission electron micrographs induced by DE and LRx. Presynaptic area is an established predictor of presynaptic origin and thalamocortical synapses are the largest excitatory synapses in layer 4 (Nahmani and Erisir, 2005). Accordingly, TEM images were collected from layer 4 (see methods; Fig. 1a), and all synaptic profiles with high contrast boundaries delineating the presynaptic membrane, active zone, postsynaptic membrane and postsynaptic density were included for analysis. Presynaptic area had the expected bimodal distribution, with a major peak at 0.24±0.007 SEM μm^2^ (presumptive IC synapses), and a minor peak at 0.688±0.2333 μm^2^ (presumptive TC synapses), which was unchanged by DE and LRx (single factor ANOVA, F = 1.4, p = 0.24, Fig. 1c). Synapses with larger presynaptic area (>0.5 μm^2^) also had larger cleft widths and PSD lengths, as expected for thalamocortical synapses (TC; Cleft width: 21.6±0.3 nm vs. 19.8±0.1 nm; PSD length: 472±30 nm vs. 356±9 nm, **p<0.01, Student’s T-test; n = 60, 189). In addition, larger synapses also contained significantly more synaptic vesicles in contact with the presynaptic membrane (presumptive “docked” vesicles) than smaller presumptive IC synapses (4.43±0.26 vs. 2.81±0.11, p<0.01, Student’s T-test) and more distance between the center of the synaptic vesicle pool and the active zone (synaptic vesicle zone distance, SVZ distance; 187±14 μm vs 147±5 μm, p<0.01, Student’s T-test; Fig 1b). This bimodal distribution of presynaptic area, with ∼15% of synapses distributed about the larger maxima, is consistent with previous estimates of TC synapse abundance (Nahmani and Erisir, 2005; Coleman et al., 2010).

**Fig. 1.**
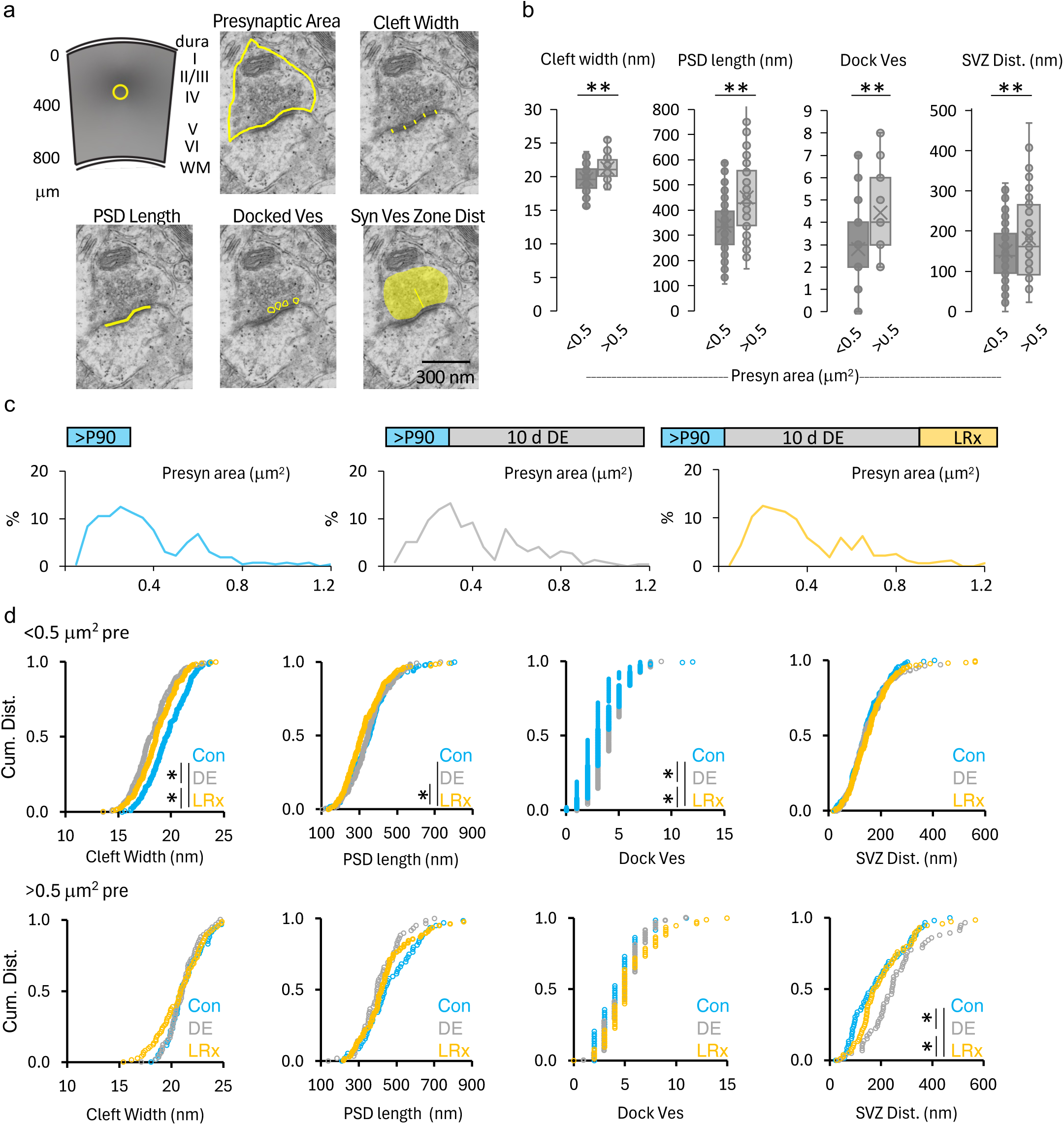
Adult DE/LRx induces excitatory synapse subtype-specific modifications in ultrastructure in mouse V1b layer 4. a) Experimental sampling area in adult mouse primary visual cortex, and representative transmission electron micrographs identifying presynaptic area, cleft width, postsynaptic density (PSD) length, docked vesicles and distance from the synaptic vesicle zone center to the active zone center (SVZ distance). b) Synaptic profiles with larger presynaptic areas (>0.5 μm^2^) have significantly larger cleft widths, PSD lengths, number of docked vesicles and SVZ distance (n = 60, 189 synapses for large and small synapses, respectively, n=5 subjects, Student’s T-test, **p<0.01). c) The distribution of presynaptic area is bimodal, and unchanged by dark exposure or light reintroduction (n = 249, 195, 294 synapses from 5, 4, 6 subjects for Con (blue), DE (grey) and LRx (yellow), respectively: One-way ANOVA, F = 1.4, p = 0.24). d) Regulation of synaptic ultrastructure by visual experience. Top: In smaller synapses (<0.5 μm^2^, presumptive intracortical) cleft width, PSD length and docked vesicle number are regulated by DE (grey) and LRx (yellow), *p<0.05, K-S test. Bottom: In larger synapses (>0.5 μm^2^, presumptive thalamocortical), only SVZ distance is significantly modified by DE (grey) and LRx (yellow), *p<0.05, K-S test.

In smaller, presumptive IC synapses, dark exposure and brief LRx (2 hr) induced widespread changes in ultrastructural parameters including cleft width, PSD length and docked vesicle number, but did not induce a change in SVZ distance (cleft width: Con = 19.6±0.14 nm, DE = 18.2±0.16 nm, LRx = 18.7±0.12 nm; PSD length: Con = 356.4± 8.6 nm, DE = 353.5±9.1 nm, LRx = 330.1±7.4 nm; docked vesicle: Con = 2.8±0.1, DE = 4.1±0.1, LRx = 3.6±0.1; SVZ distance: Con = 147±5 μm, DE = 184±18 μm, LRx = 159±6 μm; *p<0.05, n = 189, 136, 218 synapses, respectively; Fig. 1d).

In contrast, in larger, presumptive TC synapses, DE and brief LRx induced changes only in SVZ distance (SVZ distance: Con = 187±14, DE = 284±29, LRx = 207±14; cleft width: Con = 21.3± 0.2 nm, DE = 21.1±0.2 nm, LRx = 20.8±0.3 nm; PSD length: Con = 470.4± 19.5 nm, DE = 412.9±14.4 nm, LRx = 454.5±18.9 nm; docked vesicle: Con = 4.4±0.3, DE = 4.9±0.3, LRx = 5.5±0.3; *p<0.05, KS test, n = 60, 59, 76, respectively; Fig. 1d). Thus, DE and LRx induce contrasting ultrastructural changes at IC and TC synapses.

To ask if DE/LRx modified the molecular geometry of thalamic boutons, we adapted a FRET-based intermolecular proximity assay to quantify *in situ* clustering of the presynaptic scaffold protein Bassoon (Bsn) in thalamic boutons in layer 4 of adult mouse V1b (see methods and Glebov et al., 2017). A validated monoclonal anti-Bsn primary antibody was utilized, followed by anti-mouse secondary antibodies conjugated to either a FRET donor or a FRET acceptor (1:1; Fig. 2a). FRET distance is optimal at ∼10 nm, therefore and increase FRETAcceptor/Donor reports clustering of Bassoon molecules. We confirmed that the proximity assay is sensitive to intermolecular distance, as A/D_Bassoon_ decreased with serial dilution of the secondary antibody mix (500x, 1.24±0.004; 2000x, 0.95±0.003; 8000x, 0.78±0.01, n = 2343, 2304, 970 puncta, respectively. One-way ANOVA, F = 1731, p=0.0001; **p<0.01, *post hoc* Tukey, Fig. 2b). To quantify the impact of DE/LRx on presynaptic molecular organization specifically at thalamic boutons, we quantified A/D_Bassoon_ in boutons expressing vesicular transporter 2 (VGluT2; Nahmani and Erisir, 2005; Coleman et al., 2010). DE significantly decreased bouton A/D_Bassoon_, indicative of an increase in intermolecular Bsn distance. In contrast, LRx significantly increased bouton A/D_Bassoon_ indicative of a decrease in intermolecular Bsn distance (Left: Con=1.29±0.02, DE = 1.00±0.01, LRx = 1.30±0.01; n = 400 puncta/group, One-way ANOVA, F = 157, p = 0.0001; Right: Con=1.27±0.09, DE = 0.88±0.02, LRx = 1.28±0.05; n = 4, 4, 4 subjects, respectively, One-way ANOVA, F = 13.0, p = 0.002; **p<0.01, *post hoc* Tukey; Fig. 2 c, d). Neither DE nor LRx changed the total quantity of Bsn, visualized with a single secondary antibody (Left: Con = 45.4±0.2, DE = 45.2±0.1, LRx = 45.2±0.3; n = 400 puncta/group, One-way ANOVA, F = 0.25, p = 0.78; Right: Con = 45.4±1.3; DE = 43.1±1.0; LRx = 48.0±=2.5; n = 4, 4, 4 subjects, respectively; One-way ANOVA, F = 2.1, p = 0.18, Fig. 2e). The density and the area of VGluT2^+^ boutons were also unchanged by DE or LRx (Density: Con = 482±59 per 0.01 mm^2^, DE = 561±67 per 0.01 mm^2^, LRx = 503±89 per 0.01 mm^2^, One-way ANOVA, F = 0.18, p = 0.8; Area: Con = 0.58±0.03 μm^2^, DE = 0.63±0.01 μm^2^, LRx = 0.63±0.04 μm^2^, n = 4 subjects/group; One-way ANOVA, F = 0.18, p = 0.8, Fig. 2f). Thus, DE and LRx change the molecular geometry of thalamic boutons in layer 4 of V1.

**Fig. 2.**
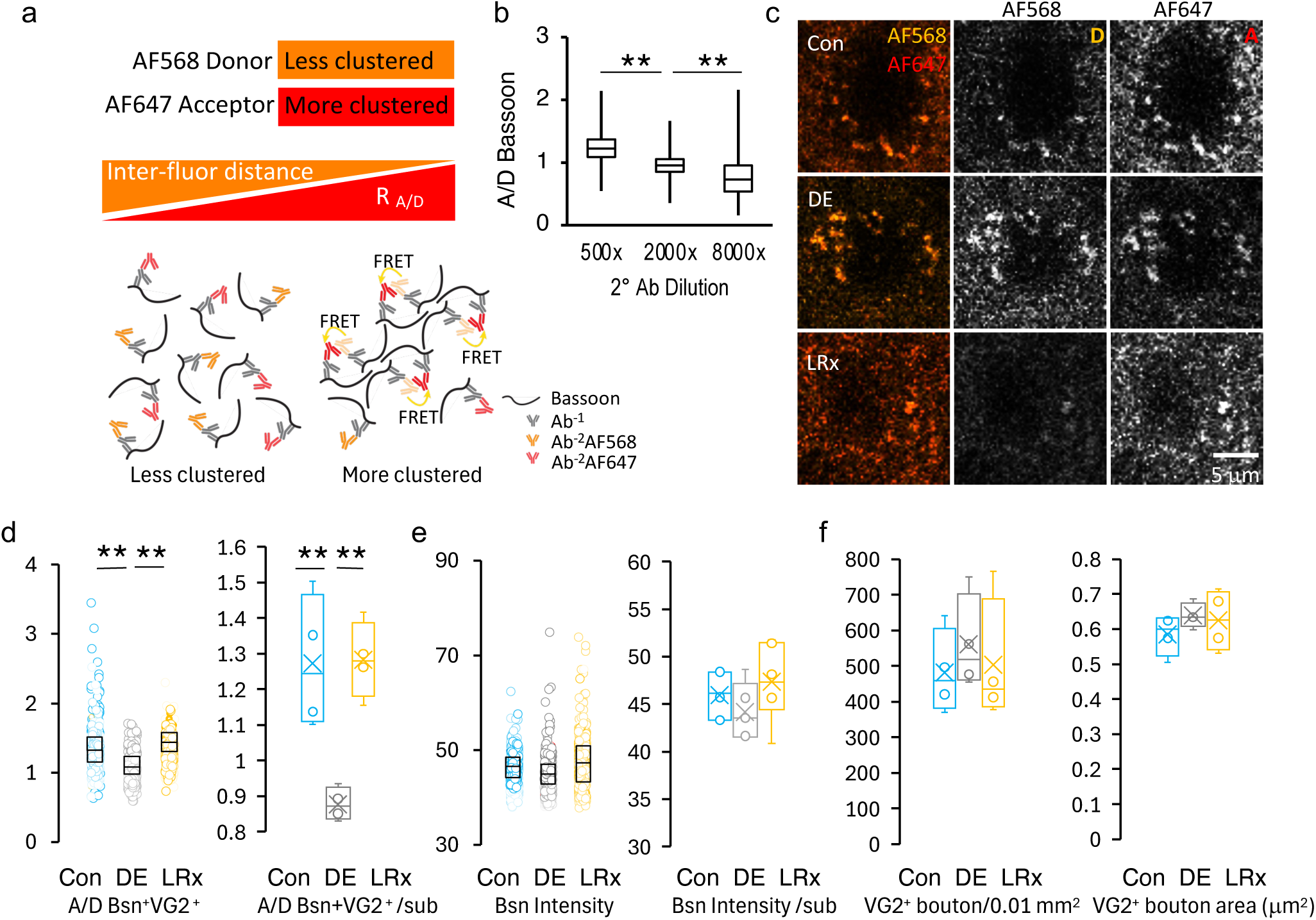
Adults DE/LRx declusters/clusters Bassoon in thalamic boutons in V1b layer 4. a) Experimental logic of FRET-based proximity assay. b) Acceptor/Donor_Bassoon_ (A/D_Bassoon_) decreased by serial dilution of secondary antibody 50:50 mix of FRET_Acceptor_ and FRET_Donor_. Box plots represent the median as a bar, 25th to 75th percentile as box, and max and min as whiskers (n = 2343, 2304, 970 puncta from 3 subjects/group; Onesway ANOVA, F = 1731, p = 0.0001; **p<0.01, *post hoc* Tukey). c) Representative examples of donor (orange, AF568) and acceptor (red, AF647) fluorescence in layer 4 of adult mouse V1b in a Con, DE and LRx subject. d) Quantification of A/D_Bassoon_colocalized with VGluT2^+^ puncta. Left: 100 puncta from each subject, subjects differentiated by color transparency, overlayed with a box plot representing the median as a bar, 25th to 75th percentile as box (4One-way ANOVA, F = 157, p = 0.0001; **p<0.01, *post hoc* Tukey). Right: Averages by subject (4 subjects/group, One-way ANOVA, F = 13.0, p = 0.002, **p<0.01, *post hoc* Tukey). e) Total Bsn immunofluorescence visualized with a single secondary antibody is unchanged by DE or LRx. Left: Bsn immunofluorescence of 100 puncta from each subject, subjects differentiated by color transparency, overlayed with a box plot that represents the median as a bar, 25th to 75th percentile as box (One-way ANOVA, F = 0.25, p = 0.78). Right: Averages by subject (4 subjects/group; One-way ANOVA, F = 2.1, p = 0.18). f) The density (left) and area (right) of VGluT2^+^ puncta in layer 4 of V1b are unchanged by DE/LRx (Density: One-way ANOVA, F = 0.18, p = 0.8; Area: One-way ANOVA, F = 0.18, p = 0.8; 4 subjects/group). Box plots represent the median as a bar, 25th to 75th percentile as box and max and min as whiskers.

Bassoon and other multi-domain scaffolds regulate synaptic vesicle organization and the density of Ca_v_ channels at the active zone. We therefore employed 2-photon (2P) live imaging to test the prediction that DE/LRx modified calcium signaling at thalamic boutons in layer 4 of V1. Axonal GCaMP (AAV5-Ef1a-DIO-Synaptophysin-GCaMP6s) was delivered to dLGN in adult VGluT2-Cre mice 3 weeks prior to recording. Axonal GCaMP imaging was performed in layer 4 of V1b at an average depth of 320 μm (Fig. 3a, b). Axonal calcium signals were evoked by 100% contrast, square wave gratings of 0.05 cpd drifting at 1 Hz delivered to the dominant eye. The inclusion criteria for visually-responsive boutons that met the following criteria were included for analysis was: 1) an evoked peak ΔF/F value (0.25 - 2.0 sec after onset of a visual stimulus in any direction) greater than 4X the standard deviation (STD) of pre-stimulus baseline (F_0_, −1.0 to 0 sec) for ≥ 40% trials and 2) the STD of ΔF/F during the rising phase of response (0.25 to 1.0 sec after visual stimulus onset) was greater than 2 X STD of baseline. DE/LRx did not change the number of visually-responsive boutons (Con = 315±34, DE = 613±289, LRx = 393±89 boutons, n = 4 subjects, One-way ANOVA, F = 0.78, p = 0.49).

**Fig. 3.**
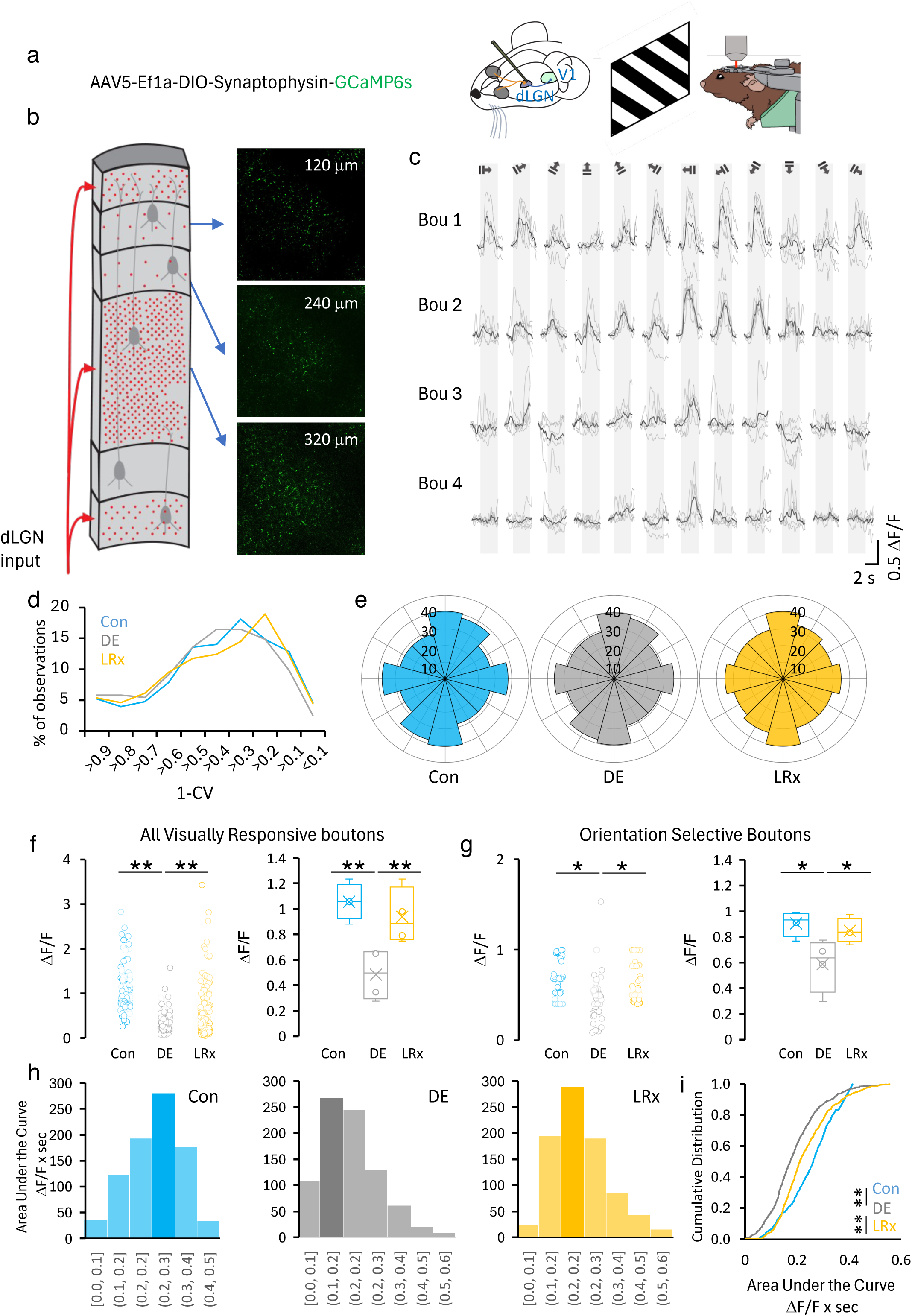
Adult DE/LRx induces reversible changes in visually-evoked and spontaneous GCaMP signals in dLGN boutons recorded in layer 4 of V1b. a) AAV5-Ef1a-DIO-Synaptophysin-GCaMP6s (750 nl) was delivered at 0.1 μl/min to dLGN: AP: 2.1 mm, ML: 2.2 mm, DV: 2.5 mm from bregma for 2P intravtial imaging of axonal GCaMP in head-fixed awake adult mice. b) Left: Depiction of the abundance of thalamic boutons throughout the vertical depth of V1b. Right: Images of axonal GCaMP6s at 3 cortical depths. c) Representative examples of visually evoked GCaMP signals in single axonal boutons (Bou) in response to 100% contrast, 0.05 cpd, square wave gratings in 12 directions drifting at 1 Hz. Grey bar = stimulus duration. d) The strength of orientation selectivity (1-CV) is unchanged by DE/LRx (all visually responsive boutons = 1258, 1225, 1572 boutons from Con, DE, LRx, 4 subjects/group; Repeated measures ANOVA between conditions, Con vs DE, F = 0.009, p = 0.93; DE vs LRx, F = 2.4x10^-6^, p = 0.99; LRx vs Con, F = 0.01, p = 0.93). e) Polar histograms of the preferred orientation of axonal boutons with 1-CV ≥ 0.33. f) Visually-evoked GCaMP ΔF/F in all visually-responsive thalamic boutons, evoked by preferred stimulus and recorded in layer 4 of adult mouse V1b. Left: ΔF/F for 30 boutons/subject, subjects differentiated by color transparency, overlayed with a box plot that represents the median as a bar, 25th to 75th percentile as box. Right: Visually evoked ΔF/F averaged by subject (1258, 1225, 1572 boutons from Con, DE, LRx, 4 subjects/group; One-way ANOVA, F = 9.9, p = 0.005, p<0.01, *p < 0.05 *post hoc* Tukey). g) Visually-evoked GCaMP ΔF/F by the preferred stimulus in axonal boutons with 1-CV ≥ 0.33. Left: ΔF/F for 30 boutons per subject, subjects differentiated by color transparency, overlayed with a box plot that represents the median as a bar, 25th to 75th percentile as box. Right: Visually evoked ΔF/F averaged by subject (n = 780, 834, 909 from Con, DE and LRx, 4 subjects/group. One-way ANOVA, F = 6.1, p = 0.02, *p < 0.05 *post hoc* Tukey). h) Spontaneous bouton calcium activity is modified by visual experience. Area under the curve for spontaneous calcium signals in 0.1 bins, largest bin highlighted. Con (blue) DE (grey) LRx (yellow). i) Cumulative distribution of AUC for all bouton calcium events shifts significantly to the left following DE and significantly to the right following LRx, (KS test, **p = 5.6 x 10^-16^, 2.2 x 10^-16^ for Con vs DE, DE vs LRx, respectively).

dLGN boutons in layer 4 were tuned for visual stimulus direction or orientation (Fig. 3c). Importantly, tuning strength, calculated as 1-CV for all visually responsive thalamic boutons was not changed by DE or LRx (Repeated measures ANOVA Con vs DE F = 0.009, p = 0.93; DE vs LRx F = 2.4x10^-6^, p = 0.99; Con vs LRx F = 0.01, p = 0.93, Fig. 3d n = 1258, 1225, 1572 boutons from 4, 4, 4 subjects for Con, DE, LRx,). Similarly, visually-tuned boutons (1-CV ≥ 0.33) had the expected preference for gratings in cardinal orientations, and this preference was unchanged following DE or LRx (Fig 3e).

In contrast, DE significantly decreased and LRx significantly increased the magnitude of visually-evoked calcium (ΔF/F) in thalamic axon boutons. Significant differences were observed across the population of all visually-responsive boutons (Con = 1.065±0.03, DE = 0.48±0.03, LRx = 0.94±0.0; n = 1258, 1225, 1572 boutons for Con, DE, and LRx, respectively, 4 subjects/group; One-way ANOVA, F = 9.9, p = 0.005), as well as the subset of thalamic boutons with 1-CV ≥ 0.33 (Con = 0.91±0.05, DE = 0.59±0.1, LRx = 0.85±0.05; n = 780, 834, 909 boutons for Con, DE, and LRx, respectively, 4 subject/groups; One-way ANOVA, F = 6.1, p = 0.02; *=P<0.05, **=p<0.01 *post hoc* Tukey; Fig. 3f, 3g). Importantly, spontaneous GCaMP signals were significantly reduced by DE and enhanced by LRx (Figs. 3h, 3i).

The bidirectional change in the thalamic bouton calcium signals by DE/LRx may reflect changes within the thalamic bouton, such as calcium channel density and distribution, or changes beyond the bouton such as a change in the excitability of dLGN neurons. As bouton calcium concentration is a primary determinant of presynaptic release probability, we employed optogenetics to quantify the impact of DE and LRx on P_r_ at thalamic boutons. TC synapses in acute slices of primary visual cortex were optogenetically isolated following delivery of channelrhodopsin (AAV5-CaMKIIa-hChR2(H134R)-mCherry) to dLGN of adult C57BL/6J mice (>postnatal day (P) 90; Fig. 4a, b). To evaluate postsynaptic changes, we measured Sr^2+^-desynchronized miniature EPSCs (Sr^2+^ mEPSCs), evoked by the half-maximal effect light intensity (470 nm, 3 msec) in the presence of TTX and 4-AP (Dodge Jr. et al., 1969; Petreanu et al., 2009; Whitt et al., 2022). In control normal-reared adult mice (NR, raised in a 12:12 h L:D cycle), light-evoked EPSCs had the expected mean amplitude of ∼13 pA, which was unchanged following 10 days of DE (Fig. 4c, d). Brief LRx (2 h) induced a small, but significant decrease in Sr^2+^ EPSC amplitudes (mean±sem: NR = 12.9±0.7 pA; DE = 13.0±0.6 pA; LRx = 11.2±0.4 pA; n = 31, 26, 25 cells from 6, 5, 5 subjects, respectively, Kruskal-Wallis H (2) = 10.39, *p <0.05 LRs v. DE, **p<0.01 LRx v Con *post hoc* Dunn’s multiple comparison test; Fig. 4c, d).

**Fig. 4.**
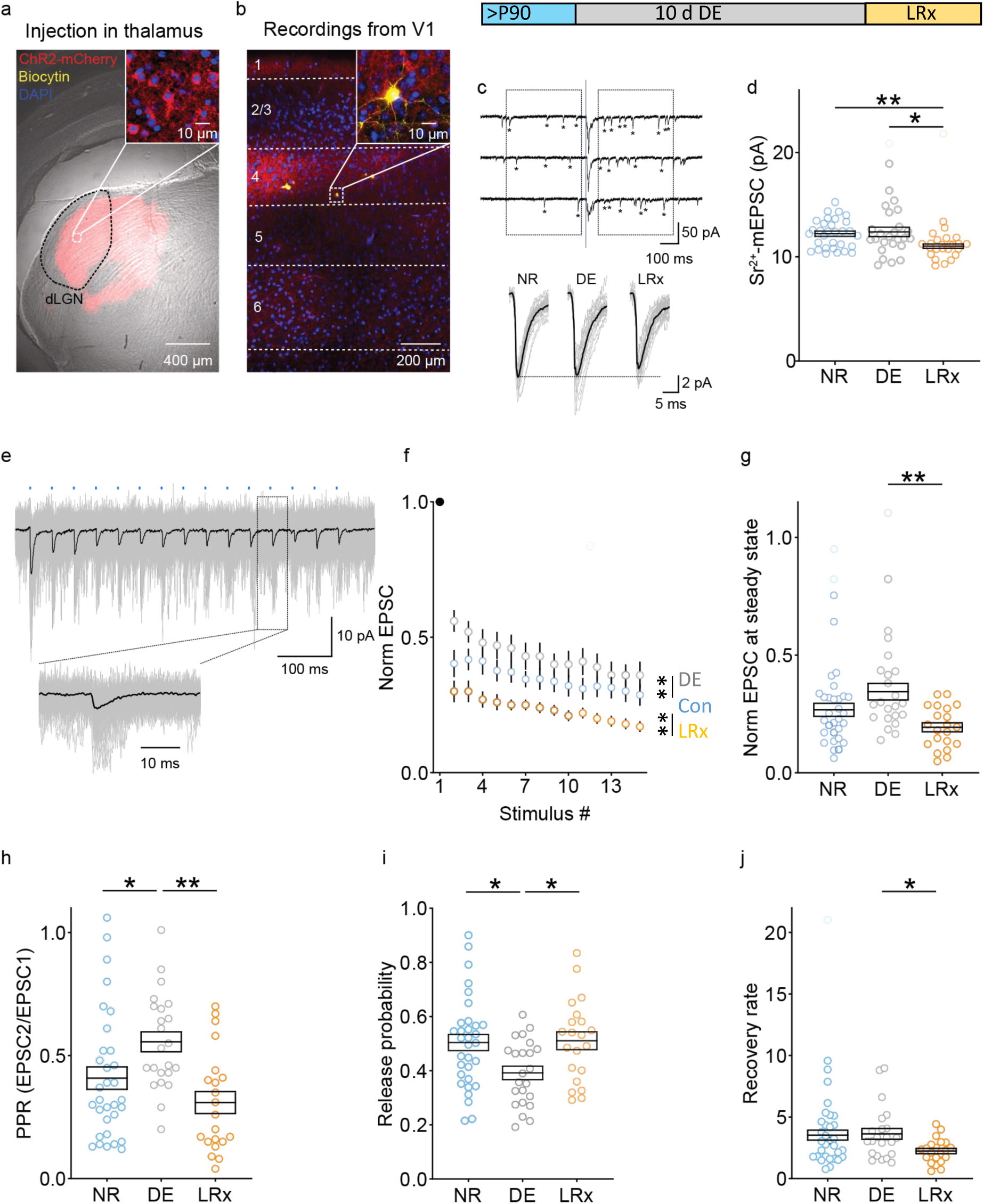
Adult DE/LRx induces reversible changes in presynaptic release probability of thalamocortical synapses in V1b layer 4. a) AAV-CaMKIIa-hChR2-mCherry delivered to the dLGN (red) effectively transfected thalamic axons. Overlay of the bright field and confocal images reveals the viral injection site in thalamus (mCherry). The inset shows dLGN neurons counterstained with DAPI (blue) and expressing ChR2-mCherry. b) Example of a pyramidal neuron in layer 4 V1b filled with biocytin during whole cell recording (yellow). Note enrichment of mCherry-positive axonal processes in L4. c) Top: Experimental timeline. Adult C57B/L6 mice (>P90) received 10 days of DE +/- LRx. Middle: Representative examples of light-evoked Sr^2+^-desynchronized miniature EPSCs (Sr^2+^- mEPSCs, 470 nm, 3 msec) in layer 4 V1b. Boxes = 400 ms collection windows for Sr^2+^-mEPSCs (asterisks) before and after LED-evoked release (blue line; 3 ms duration). Bottom: Average traces for all the cells shown in Fig. 4d (gray) and average traces for all neurons in all subjects (black) for NR, DE, and LRx. d) Sr^2+^-mEPSCs were unchanged by DE (grey) and decreased after LRx (2 hr, yellow; mean Sr^2+^-mEPSC amplitude ± sem: NR (blue), 12.9±0.7 pA; DE 13.0±0.6 pA; LRx 11.2±0.4 pA; n = 31, 26, 25 cells from 6, 5, 5 subjects, respectively, Kruskal-Wallis H (2) = 10.39, p < 0.01, *p<0.05, **p<0.01 *post hoc* Dunn’s multiple comparison test). e) Top: Repetitive optogenetic stimulation in the absence of Sr^2+^ (15 pulses of 470 nm LED for 3 msec at 25 Hz (blue dashes) at intensity that evoked a response in 100% of trials). Bottom: higher magnification of boxed recording snippet. f) EPSCs normalized to the response to the first stimulus (filled dot). Visual experience modifies the response to repetitive stimulation (n = 32, 23, 21 cells from 8, 6, 5 subjects for NR, DE, LRx, respectively. Repeated measures ANOVA, F = 63.3, **p < 0.01, *post hoc* Bonferroni). g) Normalized EPSC at steady state. h) Paired pulse ratio (EPSC in response to stimulus 2/stimulus 1) is increased by DE and recovered by LRx. i) Presynaptic release probability calculated from short-term dynamics (see Methods) is decreased by DE and increased by LRx. j) LRx decreases synaptic vesicle recovery rate. *p < 0.05, **p < 0.01 for *post hoc* tests.

In contrast, DE/LRx induced robust changes in the response to repetitive stimulation of TC synapses. Optogenetically-evoked AMPAR- dependent EPSCs were induced in layer 4 neurons by trains of LED flashes (Fig. 4e, 15 light pulses at 25 Hz at the minimal light intensity to evoke a response in 100% of trials in 100 μM APV and 20 μM bicuculline). As expected, TC EPSC amplitudes were significantly depressed by repetitive stimulation in all experimental conditions (Fig. 4f). However, the activity-dependent depression of TC EPSCs was significantly reduced by DE and increased by LRx (Two-way ANOVA: visual experience F (2, 1095) = 71.36, p<0.0001: **=p<0.01 *post hoc* Bonferroni: interaction F (28, 1095) = 0.5042, p=0.9857; n = 32, 23, 21 cells from 8, 6, 5 subjects, respectively; Fig. 4f). The normalized approximate steady state amplitude, calculated as the mean amplitude of 11^th^ to 15^th^ EPSC, was comparable to NR following DE and reduced by LRx (NR = 0.27±0.03, DE = 0.34±0.04, LRx = 0.19±0.02; n = 32, 23, 21 cells from 8, 6, 5 subjects respectively; Kruskal-Wallis H (2) = 11.22, p<001; **=p<0.01 *post hoc* Dunn’s multiple comparison test; Fig. 4g). In addition, the paired pulse ratio (PPR) significantly increased following DE and decreased following LRx (NR = 0.41±0.05, DE = 0.56±0.04, LRx = 0.31±0.04; n = 32, 23, 21 cells from 8, 6, 5 subjects respectively; Kruskal-Wallis H (2) = 13.97, p = 0.0009, *=p<0.05, **=p<0.01 *post hoc* Dunn’s multiple comparison test; Fig. 4h). Importantly, presynaptic vesicle release probability, calculated from short-term dynamics of synaptic depression in response to repetitive stimulation (see methods) was significantly decreased by DE and increased by LRx (NR = 0.50±0.03, DE = 0.39±0.02, LRx = 0.51±0.03; n = 32, 23, 21 cells from 8, 6, 5 subjects, respectively; One- way ANOVA, F = 4.699, p = 0.012, *=p<0.05 *post hoc* Tukey; Fig. 4i; as in Bridi et al. 2020 based on Wesseling and Lo, 2002). The computed recovery rate from depression was unchanged by DE and decreased by LRx (NR = 3.5±0.40, DE = 3.6±0.44, LRx = 2.2±0.22; n = 32, 23, 21 cells from 8, 6, 5 subjects respectively, Kruskal-Wallis H (2) = 6.838, p = 0.0327, *=p<0.05 *post hoc* Dunn’s multiple comparison test; Fig. 4j). Thus, synaptic plasticity at feed-forward inputs from first order visual thalamus to visual cortex is engaged by DE/LRx in adulthood and mediated by changes in presynaptic structure, organization and function.

## Discussion

Early developmental plasticity of thalamocortical synapses plays an essential role in the representation of sensory space by mapping sensory inputs to cortex, which stabilizes with age. The prediction that the loss of plasticity at TC synapses with age was irreversible was based primarily on quantification of postsynaptic variables that define Hebbian plasticity, including EPSC amplitude and slope (Crair and Malenka, 1995; Isaac et al., 1997; Feldman et al., 1998; Jiang et al., 2007). However, we found that robust and reversible plasticity can be induced at dLGN-visual cortex layer 4 synapses in adult mice by prolonged dark exposure and light reintroduction. This unexpected, experience-dependent synaptic plasticity at adult TC synapses was mediated by changes in presynaptic organization and function. DE and LRx reversibly declustered/clustered synaptic vesicles and a presynaptic scaffold protein in dLGN boutons. Visually-evoked thalamic bouton calcium signals and presynaptic release probability are reduced by DE and increased by LRx.

The presynaptic experience-dependent plasticity at TC synapses described here contrasts with the rapid, well-documented, homeostatic changes in postsynaptic response induced by DE and LRx at intracortical synapses. Interestingly, DE induces widespread changes in the ultrastructure of smaller presumptive IC synapses, including decreased cleft width, PSD size and docked vesicle number, that are consistent with a homeostatic increase in IC synaptic efficacy (Murthy et al., 2001; Lee and Kirkwood, 2019; Wen and Turrigiano, 2024; Yashiro et. al., 2005; Hengen et al., 2013; Bridi et al., 2018). In contrast, only one ultrastructural variable, the clustering of synaptic vesicles, was observed in larger, presumptive TC synapses. The thalamocortical response to DE appears to be limited to presynaptic changes in structure and function, as we observe no change in the magnitude of optogenetically-evoked TC EPSCs following DE, as previously reported (Petrus et al., 2014). A small but significant decrease in TC ESPC amplitudes was observed following LRx. We did not attempt to differentiate the potential contributions of AMPAR mobility or desensitization to this response. Our observations that DE declusters and LRx clusters synaptic vesicles support the long-standing observation that synaptic vesicle geometry is dynamic and activity-dependent. Indeed, modifications of synaptic vesicle clustering have been demonstrated following LTP induction in hippocampus and action potential suppression in cultured neurons (Glebov, et al., 2016; Harris et al., 2024) and may reflect condensation of vesicles through liquid-liquid phase separation (Milovanovic et al., 2018). As the distance of a synaptic vesicle to the active zone impacts the probability of its release, reversible, activity-dependent changes in synaptic vesicle geometry can fine-tune presynaptic function (Park et al., 2012).

The reversible declustering/clustering of synaptic vesicles at TC synapses was mirrored by declustering/clustering of the presynaptic scaffoldprotein Bassoon. Previous work demonstrates that presynaptic scaffold bassoon clusters Ca_v_ at the active zone (Davydova et al., 2014). Accordingly, DE induced both a declustering of bassoon and a decrease in the magnitude of thalamic bouton calcium signals, suggesting extensive remodeling of presynaptic geometry.

Our experiments quantified changes in the magnitude of spontaneous and visually-evoked calcium signals in dLGN boutons in visual cortex layer 4 following DE and LR. The ∼1 μm^2^ size of individual boutons and the reduced signal to noise ratio at this depth precluded longitudinal imaging. However, our population data clearly demonstrate experience-dependent modification of the Ca^2+^ signal in dLGN boutons. Importantly, we saw no differences in the strength of stimulus selectivity or the preference for cardinal orientations in the bouton population following DE or LRx. Previous work has shown that DE decreases V1 neuron response reliability was modified by DE and LRx, without a long-term impact on stimulus selectivity (Jeon et al., 2022). To ask if the magnitude of spontaneous Ca^2+^ signals at dLGN boutons was modified by DE and LRx, we quantified the area under the curve (AUC) of deconvolved calcium signals, which combine measurements of GCaMP fluorescent intensity and duration into a single value without the risk of errors introduced by spike conversions. In this analysis, we assumed discreteness and demonstrated that the AUC quanta distribution is shifted to the left following DE and shifted to the right following LRx. Although these results are consistent with a bidirectional modification in axonal calcium signaling, we cannot rule out the possibility that retinogeniculate synapses are also modified. Accordingly, monocular deprivation decreases deprived eye-evoked calcium signals in dLGN axonal boutons in visual cortex layer 1 and increases non-deprived eye-evoked spiking in dLGN (Jaepel et al., 2017; Sommeijer et al., 2017; Qin et al., 2023). In adults, the MD-induced change in dLGN neuronal responses is not impacted by silencing V1, supporting a role for retinogeniculate synapses in the plasticity of dLGN neuron spiking output (Qin et al., 2023).

Extraordinary manipulations, such as prolonged deprivation or deafferentation, may be required to engage plasticity at thalamocortical synapses in adults. For example, the strengthening of spared thalamocortical inputs contributes to the cross-modal response to nerve transection and sensory deprivation (Oberleadner et al., 2012; Yu et al., 2012; Chung et al., 2017; Jie et al., 2025; Petrus et al., 2014, 2015; Sengupta et al.,2019; Wimmer et al., 2010). Examples of unimodal strengthening of TC synapses in adults are sparse, although direct theta burst stimulation of the dLGN can induce potentiation of the field potential in cortical layer 4 (Heynen and Bear, 2001; Mainardi et al., 2010).

The high threshold for plasticity at adult TC synapses may be actively maintained by molecular brakes. Consequently, plasticity between ventral medial geniculate nucleus and the auditory cortex in adults is restored by restricting adenosine signaling from the thalamus (Blundon et al., 2017). Similarly, replacing lost plastogens, such as the neuropeptide cholecystokinin restores juvenile-like LTP at these same synapses in aged mice (Li et al., 2025). Other plastogens, such as classical psychedelics, can increase functional coupling between the thalamus and sensory networks in humans (Girn et al., 2026).

Our previous work demonstrated the activity of the matrix metalloproteinase MMP9 was obligatory for the rejuvenation of plasticity in the adult visual cortex, and that DE/LRx increased MMP9 activity at thalamocortical synapses (Murase et al., 2017). Pro-MMP9 localizes to neuronal dendrites and is released in response to synaptic stimulation (Legutko et al., 2025). Cleavage of the N-terminal pro-domain of MMP9 activity by a second proteinase activates MMP9. Future research is needed to determine if the dual activation requirements for MMP9 are met at thalamocortical synapses by the release of proMMP by hyper-active V1 pyramidal neurons by DE and the release of a pro-MMP9 targeting protease from dLGN boutons by LRx.

The age-dependent reduction in plasticity at thalamocortical synapses contributes to the developmental constraints on the reversal of amblyopia, which is achieved by dark exposure and light reintroduction. Similarly, the recovery of vision from other types of injury including stroke or retinal degeneration may be greatly facilitated by behavioral interventions that reopen the critical period of plasticity between dLGN and V1.

## Acknowledgement

This work was supported by R01EY016431 (EMQ) EY025922 (AK and EMQ), R01EY12124 (AK), and NICHD IRP (LD).

